# Detecting translational regulation by change point analysis of ribosome profiling datasets

**DOI:** 10.1101/003210

**Authors:** A. Zupanic, C. Meplan, S. N. Grellscheid, J. C. Mathers, T. B. L. Kirkwood, J. E. Hesketh, D. P. Shanley

**Author notes:** Corresponding author: Daryl Shanley, Centre for Integrated Systems Biology of Ageing & Nutrition, Institute for Ageing and Health, Newcastle University, NE4 5PL, UK.

## Abstract

Ribo-Seq maps the location of translating ribosomes on mature mRNA transcripts. While ribosome density is constant along the length of the mRNA coding region, it can be altered by translational regulatory events. In this study, we developed a method to detect translational regulation of individual mRNAs from their ribosome profiles, utilizing changes in ribosome density. We used mathematical modelling to show that changes in ribosome density should occur along the mRNA at the point of regulation. We analyzed a Ribo-Seq dataset obtained for mouse embryonic stem cells and showed that normalization by corresponding RNA-Seq can be used to improve the Ribo-Seq quality by removing bias introduced by deep-sequencing and alignment artefacts. After normalization, we applied a change point algorithm to detect changes in ribosome density present in individual mRNA ribosome profiles. Additional sequence and gene isoform information obtained from the UCSC Genome Browser allowed us to further categorize the detected changes into different mechanisms of regulation. In particular, we detected several mRNAs with known post-transcriptional regulation, e.g. premature termination for selenoprotein mRNAs and translational control of *Atf4*, but also several more mRNAs with hitherto unknown translational regulation. Additionally, our approach proved useful for identification of new gene isoforms.

## Introduction

Recent studies have shown that only 30-60 % of the variation in cellular protein levels can be attributed to mRNA levels (Tian et al. 2004; Vogel et al. 2010). The remainder is most likely due to post-transcriptional regulation, including mRNA transcript-specific regulation of translation, and protein degradation (Cho et al. 2006; Brockmann et al. 2007; Sutton et al. 2007; Sonenberg and Hinnebusch 2009; Rogers et al. 2011). While translational regulation occurs largely during initiation (Richter and Sonenberg 2005), recent experimental and computational studies have discovered that regulation of elongation also plays an important role, by means of cis-regulatory elements on the transcripts, codon bias or mRNA structure (Arava et al. 2005; Dittmar et al. 2006; Kudla et al. 2009; Tuller et al. 2010; Leprivier et al. 2013).

Ribosome profiling (Ribo-Seq), a technique based on deep-sequencing of mRNA regions protected by ribosomes, which provides the positions of ribosomes on the entire transcriptome (ribosome profiles), has enabled the study of translation regulation at the genome-wide level (Ingolia et al. 2009; Ingolia et al. 2012). Analysis of Ribo-Seq datasets using novel bioinformatic algorithms led to the identification of cis-regulatory elements that control the speed of elongation (Ingolia et al. 2011; Stadler and Fire 2011; Li et al. 2012), the sequence of events in miRNA regulation of translation (Guo et al. 2010; Bazzini et al. 2012) and new translation initiation sites (TIS) (Ingolia et al. 2011; Lee et al. 2012). The full scope of ribosome profiling studies is presented in a recent review (Ingolia, 2014). A recent study of proteotoxic stress revealed a decrease in ribosome density on ribosome profiles of affected mRNA transcripts (Liu et al. 2013). However, despite biological relevance, to date no systematic genome-wide search for changes in ribosome density has been performed.

The aim of the present work was to develop a computational approach for detecting translational regulation, especially regulation during elongation, from ribosome profiles. We started from the assumption that most translational regulation events (e.g. ribosome stalling, alternative termination or alternative initiation) cause changes in ribosome density along the translated mRNAs. After determining by mathematical modelling that it is theoretically possible to discriminate between different forms of translational regulation based on the patterns of ribosome density, we used a change point algorithm (Erdman and Emerson 2008) to find changes in ribosome density within ribosome profiles using Ribo-Seq data generated for mouse embryonic stem cells (mESCs) (Ingolia et al. 2011). This analysis detected many mRNAs with known translational regulation, as well as new translational regulation targets of alternative termination, alternative initiation and ribosome drop-off. In addition, the analysis revealed the presence of several new gene isoforms.

## Results

### Simulation of protein synthesis and translational regulation

To analyze the effects of translational regulation on protein synthesis and ribosome profiles, we built a family of totally asymmetric simple exclusion process (TASEP) models of protein translation (Figure 1, Supplementary File1) (Lakatos and Chou 2003; Shaw et al. 2003). The models include two species: mRNA molecules with *N* codons and (an infinite pool of) ribosomes of size *L* codons. Free ribosomes bind to the available initiation codon of the mRNA at rate *k*_*I*_, before progressing along the mRNA at a codon specific rate *k*_*Ei*_(*i = 1,2,…, N*), and leaving the mRNA at the last codon at rate *k*_*T*_. Each ribosome is allowed to progress only if there is no steric hindrance from the ribosome in front. Additional reactions, some of which have not been addressed previously with TASEP models, include: scanning past the canonical TIS to an alternative TIS in the coding region of the mRNA (CDS) at rate *k*_*Alt*_ and ribosome drop-off or alternative termination at rate *k*_*Di*_.

**Figure 1.**
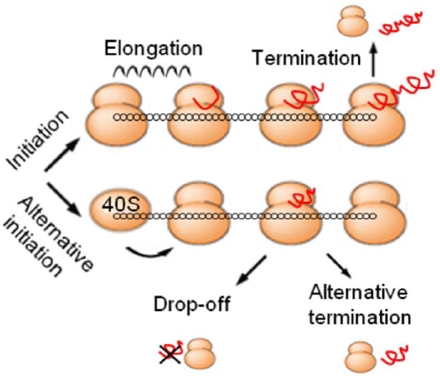
TASEP model of translation. Model reactions include: canonical translation, i.e. initiation by free ribosomes at the canonical TIS, codon specific elongation and termination at canonical termination site (with production of full-size peptide), and additional translational regulation events, i.e. alternative initiation downstream of the canonical TIS, ribosome drop-off and alternative termination.

Depending on initiation, termination and (codon-specific) elongation rates, different regimes of elongation are possible, which result in significantly different protein synthesis rates and different ribosome profiles (Lakatos and Chou 2003; Shaw et al. 2003). If, at first, we assume the elongation rate is codon independent, then at low initiation rates the mRNA is sparsely populated by ribosomes, and the protein synthesis rate is low. Increasing initiation rate increases ribosome density and protein synthesis, until steric hindrance between ribosomes prevents a further increase (Figure 2A, Figure S1A). Conversely, slow termination leads to low protein synthesis as a result of ribosome queuing; increasing termination eliminates the queues. Any further increase in termination does not change the density of protein synthesis (Figure 2B, Figure S1A). The oscillations apparent for low *k*_T_ are due to a queue of ribosome spanning the length of the mRNA (Figure 2B).

**Figure 2.**
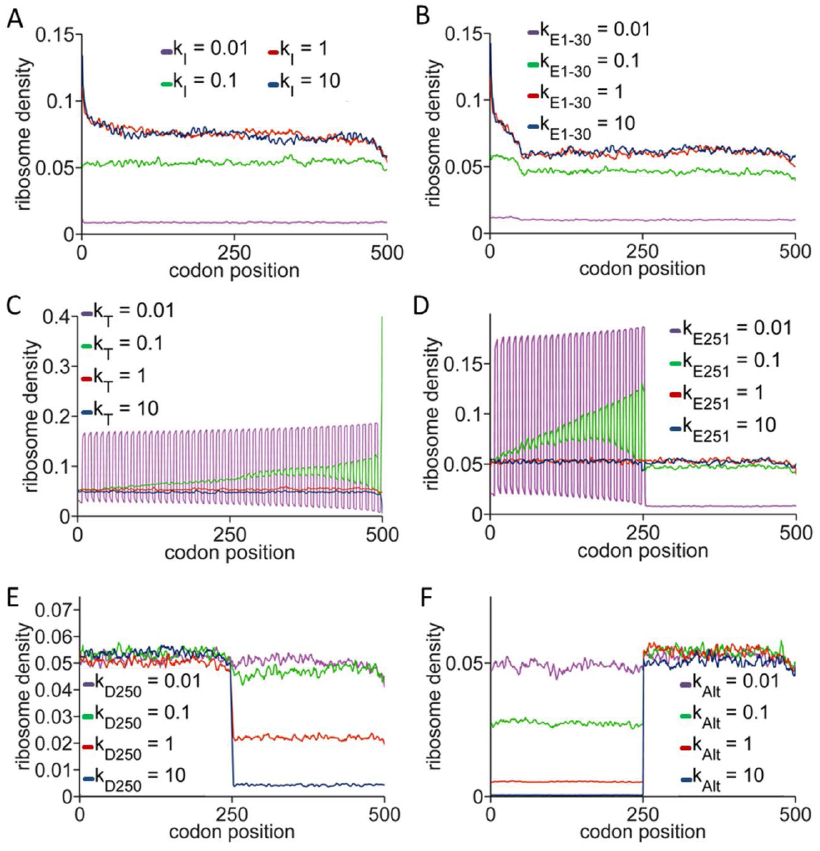
TASEP simulations of translational regulation. Ribosome density along the mRNA coding region for different rates of (A) canonical initiation, (B) elongation of the first 30 codons – ramp, (C) termination, (D) elongation of a central codon, (E) ribosome drop-off, (F) alternative initiation downstream of canonical TIS. Unless stated otherwise, *k*_*Ei*_ *= 1*, *k*_*I*_ *= 0.1*, *k*_*T*_ *= 1*, *k*_*Alt*_ *= 0*, *k*_*Di*_ *= 0*. For all simulations, *N = 500*, *L = 10*, *n = 10000* (number of samples taken from the simulation).

If the elongation rate is codon dependent, there are several possible scenarios. Slow codons at 5’ positions have effects similar to slow initiation and slow codons at 3’ positions effects similar to slow termination (Figure S2). Slow codons in the central region of the CDS, with elongation rates higher than the initiation rate, cause a peak in the ribosome profile at their respective positions; lower elongation rates cause a queue behind the stalled ribosome, with few ribosomes in front; finally at very low elongation rates the queue reaches to the first codon and begins to limit initiation (Figure 2C).

A special case of putative regulation of ribosome progression along the mRNA is the “ramp”, a stretch of the first 30-50 CDS codons that are more likely to include rare codons, which was discovered by ribosome profiling in yeast and further investigated in a computational study of codon bias (Ingolia et al. 2009; Tuller et al. 2010). Decreased elongation rates of the first 30 codons are associated with increased ribosome density (Figure 2D). Nevertheless, since similar increased ribosome density is also predicted when initiation rates are high and elongation rates normal (compare Figures 2A and 2D), rare codons may not in fact be the cause of the ramp. Alternatively, the “high initiation rate” ramp may be caused by boundary effects of the finite TASEP and occurs if translation is occurring at close maximal coverage of the mRNA with ribosomes. Since the ramp associated with rare codons can occur at any coverage, we propose that measuring ribosome coverage could be used to differentiate between the two possible natures of the ramp and the corresponding mechanisms.

We also found that all other simulated cases of translation regulation had significant effects on ribosome density, indicating that the analysis of changes in ribosome density could indeed lead to identification of translationally regulated mRNAs. Alternative initiation results in a step-like increase in density, while ribosome drop-off and alternative termination result in a step-like decrease (Figure 2E-F; Figure S2C).

### Change point algorithm robustly detects translation regulation from *in silico* generated data

After performing the simulations, we tested whether the change point algorithm could detect the changes in ribosome density in computationally generated ribosome profiles. In brief, the algorithm detects points along the length of the mRNA where the ribosome density is changed, while ignoring the effects of noise (see Methods). If the change is abrupt, the change point (CP) position can be determined very exactly, while for slower changes one or more CPs may be detected, with larger positional uncertainty. Although the algorithm can detect very small changes, in this work we limited the search to only those changes that were larger than 33 % and so only CPs most likely to include strong translation regulation were included in the analysis.

The CP algorithm provided a robust method for detecting changes in density for all the translational regulation modes that we tested. A single slow elongation codon, slow termination, alternative initiation and ribosome drop-off/alternative termination all led to step-like changes in ribosome densities and were also detected as such. Slow termination and slow elongation that led to ribosome stalling and queues were detected as oscillations in ribosome density (with the caveat that, for detecting ribosome queues, different algorithm parameters had to be used - see Methods). The minimum/maximum parameters at which change point detection was robust are presented in Table S1, with example outputs from the CP analysis in Figure S3.

### Bias in ribosome profiling

The next step was to use the CP algorithm to detect translation regulation events in a Ribo-Seq dataset. We chose a recent mouse embryonic stem cell study (Ingolia et al. 2011) because it included both Ribo-Seq and RNA-Seq datasets obtained from the same cell culture. A recent analysis of a different Ribo-Seq dataset from the same study indicated increased ribosome flux along the translated mRNAs (Dana and Tuller 2012). This implied that more ribosomes translated the 3’ regions of the CDSs than the 5’ regions, a feature which is inconsistent with current biophysical models of translation. Since such a ribosome profiling bias would likely result in a high number of false positives detected by the CP algorithm, we decided to search first for potential bias in our dataset.

From the combined Ribo-Seq/RNA-Seq dataset, 8,933 well-expressed mRNA transcripts were selected with high enough signals in both Ribo-Seq and RNA-Seq (Table S2). The averaged ribosome and RNA-Seq profiles of the 8,933 highly-expressed mRNAs confirmed the presence of bias in the datasets. The ribosome density was highest at initiation and termination codons and then dropped towards the middle of the CDS. However, unexpectedly, the central region of the CDS exhibited higher ribosomal density than the 5’ and 3’ ends (Figure 3A). It should be noted that this bias towards high ribosome density in the central region of the CDS is much more apparent if the analyzed mRNAs are normalized according to the length of their CDS before averaging ribosome density, rather than if ribosome density is averaged for the first and last 50 codons, as was done in the original Ribo-Seq study (compare Figure 3A and Figure S4) (Ingolia et al. 2011). Importantly, the average ribosome profile matched the average RNA-Seq profile very well, suggesting that the ribosome profiling bias is caused by a factor common to Ribo-Seq and RNA-Seq. Indeed, dividing the average ribosome profile with the average RNA-Seq profile results in a ribo/RNA average profile that is consistent with current biophysical models of translation: ribosome density decreases slowly from the initiation to the termination codon (Figure 3A).

**Figure 3.**
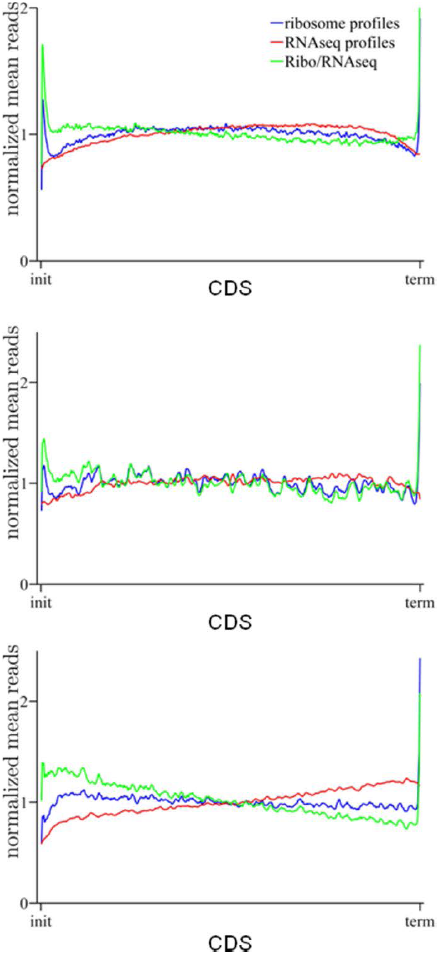
Average ribosome, RNA-Seq and normalized profiles. Average read densities for Ribo-seq, RNA-Seq and ribo/RNA for (A) 8,933 well expressed mRNAs, (B) 342 single exon mRNAs and (C) 630 long mRNAs. All read densities are normalized against the mean number of reads of the profiles and against their length.

Additionally, analyzing individual ribosome (and RNA-Seq) profiles revealed that ribosome (and RNA-Seq) density along the transcripts was very rarely constant, as would be expected for translation without strong regulation. Instead, many step-like changes were present in the profile. Two likely reasons were alignment of RNA reads to multiple genomic regions and assignment of RNA reads aligned to single genomic regions to multiple gene isoforms that result from alternative splicing, which have both featured in the original study (Ingolia et al. 2011) (Figure 4). The profile of a single mRNA transcript can thus include not only the counts of its own reads, but also the counts of reads belonging to all alternatively spliced isoforms that share the same sequence.

**Figure 4.**
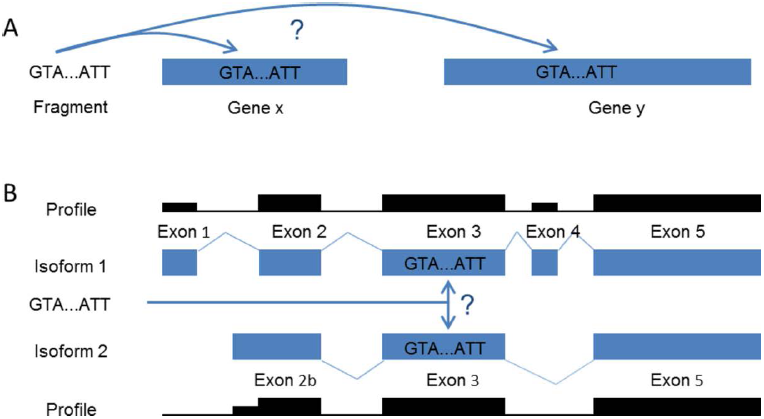
Multiple alignments and multiple assignments of RNA reads. (A) If an RNA read matches more than one genomic sequence it can be assigned to both positions. (B) If there are more annotated isoforms that share the same genomic sequence, RNA reads can be assigned to all isoforms, leading to step-like changes in ribosome and RNA-Seq profiles.

Because the effects of multiple alignments were estimated to be small in previous studies (Ingolia et al. 2011; Dana and Tuller 2012), we decided to focus on multiple assignments. We limited the dataset to a group of single-exon genes without known isoforms, for which any multiple assignment should not be possible. This decreased the bias but did not eliminate it (Figure 3B). Interestingly, choosing only long mRNAs, as was done in the study that found the ribosome profiling bias (Dana and Tuller 2012), also did not eliminate the bias, but this choice changed the average ribosome density: the RNA-Seq profiles for long mRNAs increase from the initiation to the termination codon and consequently the ribo/RNA profiles fall sharply in the same direction (Figure 3C). This indicates that the Ribo-Seq and RNA-Seq bias depend on the length of the transcript.

### Change point detection in ribosome and RNA-seq profiles

Although multiple assignments could not completely explain the Ribo-Seq bias that was seen from the average profiles, they caused undesirable step-like changes in ribosome profiles and needed to be removed before the CP analysis. Therefore, assuming that approximately the same number of multiple assignments and multiple alignments occurred in both Ribo-Seq and RNA-Seq, all individual ribosome profiles were normalised with the corresponding RNA-Seq profiles. In this way, CP analysis of ribosome and RNA-Seq profiles should detect alternative splicing and translational regulation, while the analysis of the derived ribo/RNA profiles should detect translational regulation only.

The CP algorithm was run for 8,933 ribosome profiles, RNA-Seq profiles and ribo/RNA profiles using the same parameter values. As expected, the algorithm detected the fewest CPs in the ribo/RNA profiles (Ribo-Seq: 15483 CPs, RNA-Seq: 22342 CPs, ribo/RNA: 8255 CPs) (Table S2). Almost half of the CPs that were found in ribosome profiles, but no longer present in ribo/RNA profiles, matched the positions of known exon junctions (downloaded from UCSC Genome Browser database, assembly NCBI37/mm9; Table S1) (Karolchik et al. 2004). The necessity of the normalization approach was also evident when examining individual transcripts; for example, nine CPs were found in both ribosome and RNA-Seq profiles for ribosome biogenesis regulator (*Rrs1*) and were all eliminated by the normalization, while for glutathione peroxidase 1 (*Gpx1*) two CPs were found in both ribosome and RNA-Seq profiles and one CP remained after normalization (later identified as a true translation regulation event (Figure 5)). Nevertheless, when examining positions of CPs found in all three types of profile, it emerged that normalization not only removed CPs, but also added new ones, particularly in the 5’ and 3’ ends of the CDSs (Figure S5). As most of these extra CPs were most likely to be artefacts of normalization, we took particular care to remove them in subsequent analysis.

**Figure 5.**
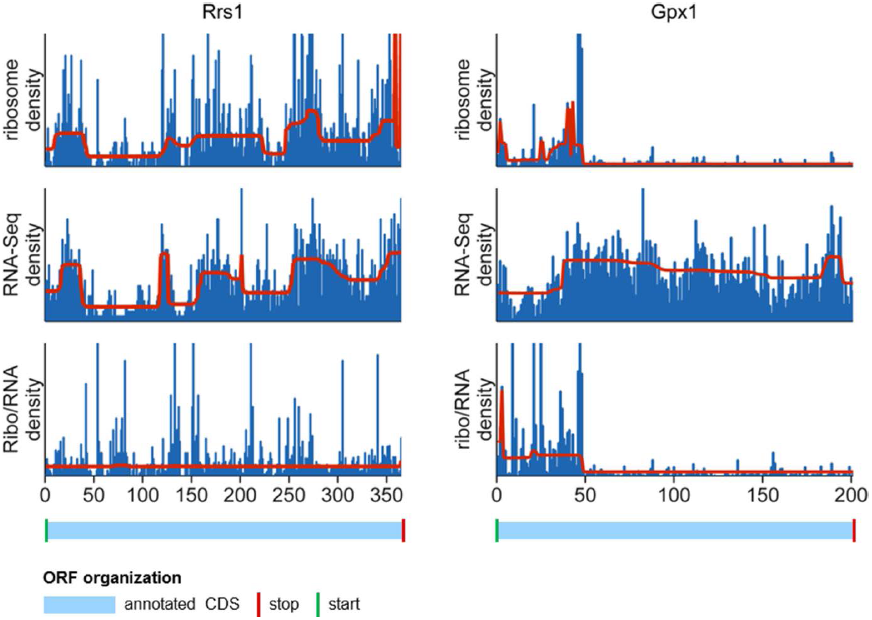
Elimination of artefacts from ribosome profiling. RNA read densities are shown in blue and the segments of equal density estimated by the change point algorithm are shown in red. After normalization by RNA-Seq, artefacts, such as effects of multiple alignments and assignments, are removed from ribosome profiles. (A) For *Rrs1* normalization removed all artefacts from the ribosome profile, consequently the change point analysis of the normalized profile resulted in no CPs found. (B) For *Gpx1*, a single change point remained after normalization, which we later showed to be a true translation regulation event. The annotated CDSs are shown below the density plots in light blue. Green vertical lines indicate annotated start codons, red lines indicate annotated stop codons, and black lines indicate exon-exon junctions.

In total, 34,844 CPs were found in all the different profiles. To find mRNA targets with a high probability of translation regulation, we applied several filters to this CP set. First, we categorized the CPs based on the profile (ribosome, RNA-Seq or ribo/RNA) in which they were found (Figure S6): translational regulation targets should have CPs in ribosome profiles, but not in RNA-Seq profiles; and normalization of the ribosome profiles with RNA-Seq profiles should not eliminate the CP. As a second filter, we eliminated all those CPs that coincided with known exon junctions. We were left with 815 CPs belonging to 635 mRNAs. From these, 336 CPs that came in pairs (an increase in signal density followed by a decrease to the same level, or *vice versa* - see exon 4 in Figure 4) were removed as this pattern strongly suggested alternative splicing coupled with multiple assignment of RNA reads (see Figure 4). Gene ontology analysis of the remaining translation regulation candidates (479 CPs on 462 mRNAs) showed enrichment of several biological processes and cellular components, including nucleobase-containing compound metabolic process (GO:0006139), translational initiation (GO:0006413) and nuclear part (GO:0044428) (for a full list see Table S2).

### Classification of CPs

Using a decision tree, the CPs were then classified into the following groups: alternative termination, alternative initiation, ramp, drop-off, slow termination, stalling and false positives (Figure S7). After automatic classification, each mRNA profile was visually inspected and compared with the genomic information available in UCSC Genome Browser (Meyer et al. 2013). If the CP could be explained by a factor different from translational regulation (e.g. by alternative splicing/mutiple assigment), it was regarded as a false positive.

This analysis discovered six genes with potential for strong alternative termination: *Gpx4*, *Tmem55b*, *Atf4*, *2700094K13Rik*, (also known as SelH), *Sep15* and *Gpx1* (Figure 6, Table S2). For all six genes, the CPs are positioned at internal stop codons which are distant from exon-exon junctions, indicating they are not artefacts of multiple assignments. Four of the six correspond to selenoprotein genes (*Gpx4*, *SelH*, *Sep15* and *Gpx1*) with a UGA codon in the central region of the CDS. The presence of a stop codon in the CDS is a characteristic feature of seleonoprotein mRNAs where it codes for the amino acid selenocysteine (Hesketh 2008). There are 24 known mouse selenoprotein genes but this CP analysis did not detect the other 20 genes. Likely reasons for this finding include: only six of the other 20 selenoprotein genes were well expressed and thus part of our analysis, with four of the six (*Txnrd1*, *SelK*, *SelT*, and *Vimp* (also known as *SelS*) having the selenocysteine codon positioned immediately before the stop codon making detection of alternative termination impossible. In addition, for the remaining two genes (*Sepw1*, *Msrb1 (*also known as *Sepx1)*), CPs were found in ribosome profiles, but then lost during normalization (*Sepx1*) or because the UGA was near an exon junction (*Sepw1*) (Figure S8).

**Figure 6.**
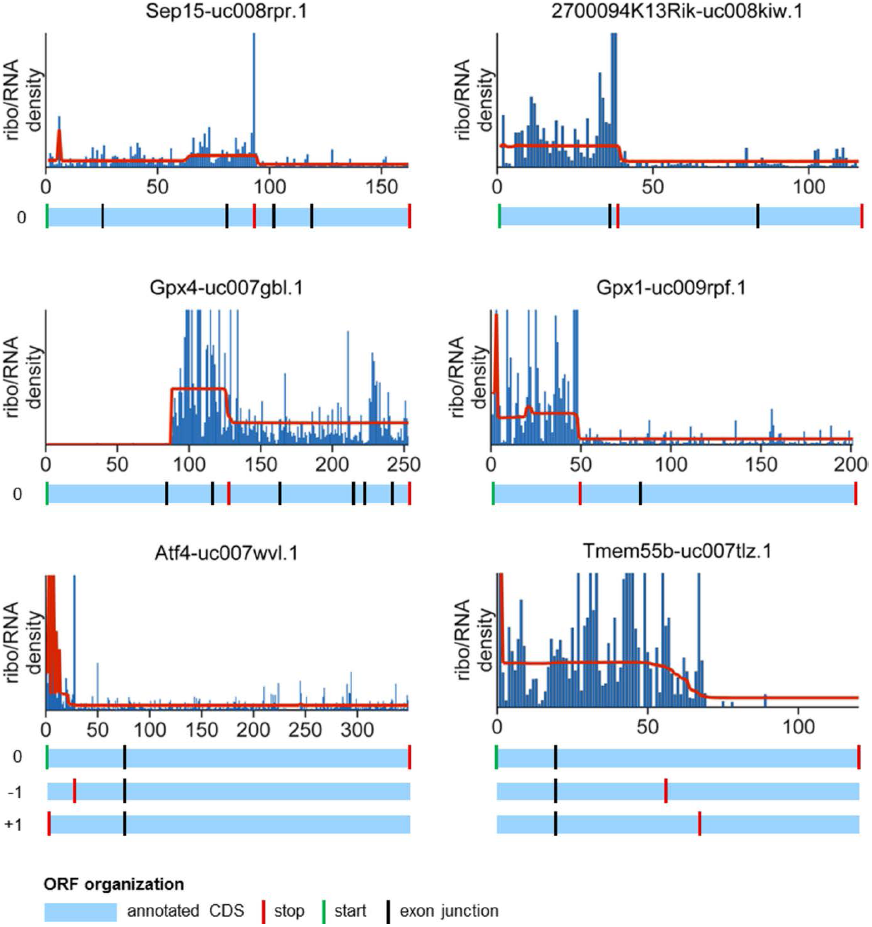
Genes with alternative termination. Ribo/RNA profiles are shown in blue and the segments of equal density estimated by the change point algorithm are shown in red. The annotated CDSs are shown below the profiles plots in light blue. For *Atf4* and *Tmem55b* the three reading frames are indicated as 0,-1 and +1.Green vertical lines indicate annotated start codons, red lines indicate annotated stop codons and/or the first stop codon encountered by the ribosome, and black lines indicate exon-exon junctions.

In contrast to the mRNAs from selenoproteins which were translated in frame 0, the translation of the remaining two mRNAs (*Atf4* and *Tmem55b*) was frame-shifted. For *Atf4,* we found two functional TISs, the canonical one and one in the 5’UTR in frame -1, indicating high translation of an uORF. For *Tmem55b,* we discovered a functional TIS in the 5’UTR in frame +1, while the canonical one was not functional (Figure S9). For both proteins, the translation was terminated in the middle of the annotated CDS, with potential production of functional peptides.

We also discovered 11 genes with potential alternative initiation in the CDS leading to substantial translation: *Lactb2*, *Samm50*, *Mrpl24*, *Cyr61*, *Ube2j2*, *Psmc2*, *Psmd8*, *Lsm14a*, *Nsmce1*, *Mylpf* and *Echs1* (Figure 7, Table S2). Of these, five genes (*Ube2j2*, *Psmc2*, *Psmd8*, *Nsmce*, *Mylpf*) were also discovered by Ingolia et al when initiation was stopped with the translation inhibitor Harringtonine (Ingolia et al. 2011). Most of the discovered mRNAs were expressed in frame 0 in their annotated form; however due to two functional TIS, the annotated and an alternative peptide were produced. *Psmd8* (and possibly *Psmc2*) was expressed as an unknown isoform (Figure 7).

**Figure 7.**
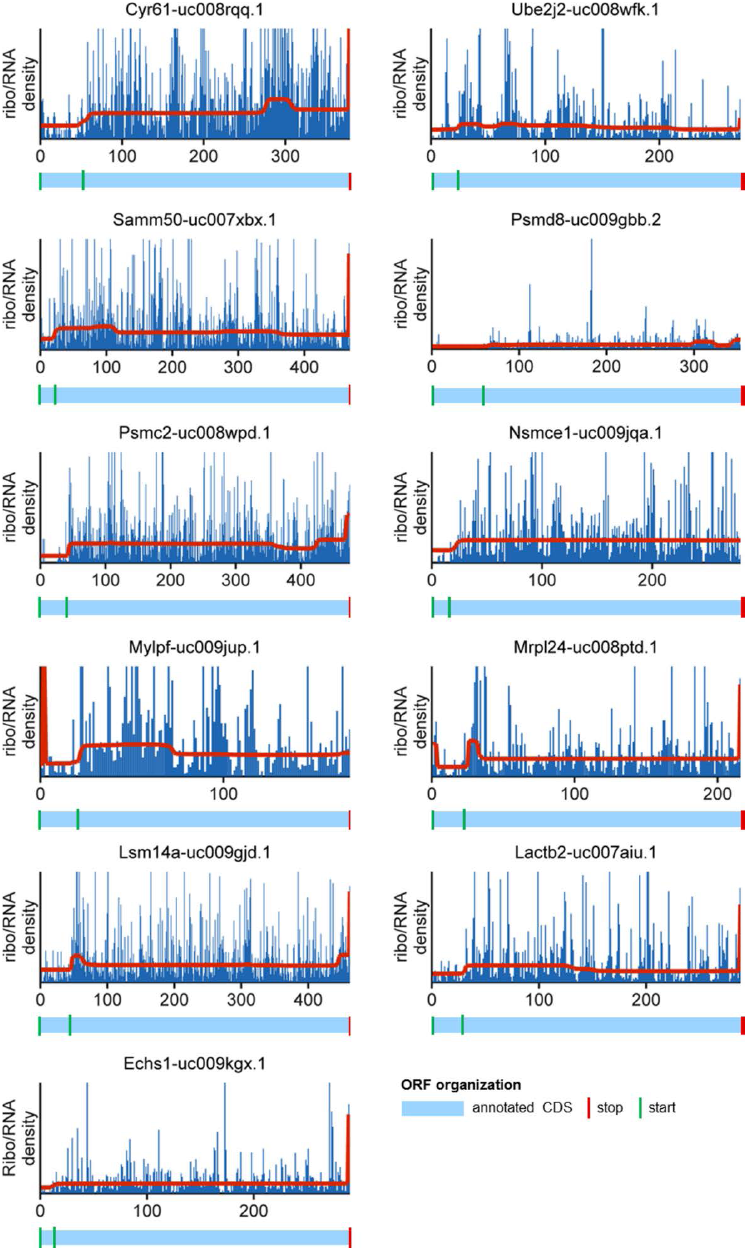
Genes with alternative initiation. Ribo/RNA profiles are shown in blue and the segments of equal density estimated by the change point algorithm are shown in red. The annotated CDSs are shown below the profiles plots in light blue. Green vertical lines indicate annotated start codons and the newly discovered alternative start codons, while red lines indicate the annotated stop codons.

We then tested whether CP analysis was useful for establishing the level of translation from alternative initiation sites predicted by before (Ingolia et al. 2011). We compared all 3,426 alternative TISs predicted by Ingolia et al. to the positions of the detected CPs in ribosome profiles and found 13 matches (Figure S10; *D10Jhu81e, Laptm4a, Enox1, Eif4g1, Etv5, Hn1l, Ppp1ca, Hnrnpul2, Hnrpa3, Nrd1, Mtf2, Isg20l1*, *Flna*). Of these, most associated open reading frames (ORF) were in the canonical frame and produced truncated proteins, while *Mtf2*, *Isg20l1* and *Ppp1ca* were translated in frame +1 and produced a short peptide only, which can be seen by the decrease in ribosome density shortly after the alternative initiation/increase in density (Figure S9).

The analysis found eight genes with potential ribosome drop-off (*Pum2, Cinp, Spin1, Kat6b, L3mbtl2, Uhrf1, Zmat3,*and *Pde6g*) (Figure S10, Table S2). In most cases, the drop in ribosome density was preceded by a large spike, indicating the drop could be a consequence of stalling at a slow codon. The drop-off group was enriched for methylated histone residue binding (GO: 0035064) (*Spin1, L3mbtl2, Uhrf1*; FDR = 0.017), indicating a possible role for particularly slow elongation in the correct folding of proteins that bind histone residues. However due to small sample size, this interpretation has to be taken with due caution. Interestingly, the genomic sequence of the eight genes around the predicted drop-off was enriched for G and A nucleotides, in particular the gaa codon (Figure S11).

We found 97 mRNAs with the ramp - high ribosome density immediately after initiation (Table S2). When this group was compared with all well-expressed genes for translational efficiency no difference was found, indicating that slow elongation at the start of the CDS and not fast initiation, which was a competing hypothesis broad forward by the modelling, is the main cause of the ramp (Figure S12). We also found no enrichment for rare codons or for any gene ontology (GO) terms. Slow termination was predicted for 45 mRNAs (Table S2). We found no enrichment for any particular stop codon, unusual termination context, rare codons preceding the stop codon or any GO terms for this group.

Despite efforts to eliminate false positives, approximately 60 % of CPs on the final list could not be attributed to the tested translational regulation mechanisms. Detailed manual analysis showed that many CPs were a consequence of alternative splicing; however because the CPs found around the known splice sites were eliminated, only CPs at unknown splice sites should have remained. By comparing the whole set of 462 candidates for translational regulation with UCSC isoform information, we found new isoforms for 31 genes (Figure 8, Table S2), suggesting that CPs could also be good markers for isoform discovery.

**Figure 8.**
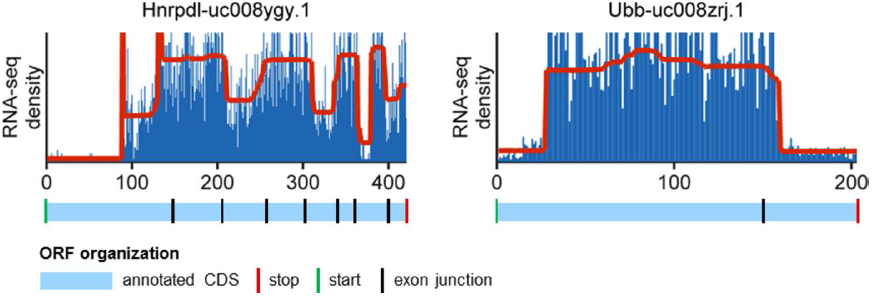
Genes with new isoforms. RNA-seq profiles are shown in blue and the segments of equal density estimated by the change point algorithm are shown in red for *Hnrdpl* (left) and *Ubb* (right). The annotated CDS are shown below the profiles in light blue. Green vertical lines indicate annotated start codons and red vertical lines annotated stop codons, while black vertical line indicate exon-exon junctions. For both genes the change point location differs significantly from the annotated exon-exon junctions and indicates hitherto unknown spicing sites. The remaining new isoforms found are described in Table S2.

## Discussion

In this study we demonstrated how detecting changes in ribosome density along mRNA molecules can be used to find targets of translational regulation. By focusing on large changes of ribosome density we were able to identify mRNA molecules for which translational regulation determines the sequence and number of expressed proteins. This approach detected known, as well as unknown, targets of alternative termination, alternative initiation and putative ribosome drop-off. In addition, during our analysis we determined that Ribo-Seq suffers from the same experimental bias as RNA-Seq and showed the utility and necessity of analysing the results of both methods in parallel. Finally, we also demonstrated that changes in ribosome density or RNA-Seq density can be used to detect new genetic isoforms.

Several studies have used Ribo-Seq to study translational regulation after different interventions, and most have used translational efficiency of groups of transcripts as their measure of interest (Ingolia et al. 2011; Bazzini et al. 2012). One study that used individual ribosome profiles and position-based ribosome density in its analysis, came to the surprising conclusion that ribosome flux increased along the transcripts with more ribosomes present at the 3’ part of the CDS than the 5’ part (Dana and Tuller 2012). The authors offered two explanations for this proposition, either extensive alternative initiation from TISs inside the CDS or an unknown bias present in the dataset. Our analysis of the Ribo-Seq and RNA-Seq datasets identified a bias and provided an explanation. The average ribosome and RNA-Seq profiles had highest ribosome density in the middle of the CDS, with lower density at both the 5’ and 3’ ends (Figure 2). When the ribosome profiles were normalized by corresponding RNA-Seq profiles, the unusual density distribution disappeared and the resulting ribo/RNA profiles had a very small decrease from the 5’ to the 3’ region. This is consistent with the current understanding of translation - a small fraction of ribosomes drop-off the mRNA transcripts and do not finish translation (Kurland 1992). We repeated the analysis using only long transcripts to better match Dana and Tuller’s analysis. This time the average Ribo-Seq and RNA-Seq were not so similar, but normalization still led to the expected ribo/RNA profiles. The sharper decline in ribo/RNA normalized profiles for longer mRNA molecules (compare Figure 2A and 2C) is not surprising. If the slow decrease in ribosome density from 5’UTR to 3’UTR seen in ribo/RNA normalized profiles of all genes (Figures 2A-B) is due to a low probability of ribosome dropping off the transcript, then the faster decrease in the normalized profiles of longer mRNAs can be explained by their length alone. Our findings suggest that the increased ribosome flux reported by Dana and Tuller could be an artefact of the Ribo-Seq and RNA-Seq protocol used in the original study.

One possible cause of the observed bias could be assignments of RNA reads to multiple genetic isoforms (Figure 3). To test this hypothesis, the same analysis was run on only single-exon transcripts. This eliminated part, but not all, of the bias. Another possible source of the bias is the so called fragment bias (Bohnert and Ratsch 2010; Hansen et al. 2010; Roberts et al. 2011) which has been shown to depend on transcript length and could therefore also explain the observed differences in profiles between long and short transcripts. Apart from being a source of part of the Ribo-Seq bias, multiple assignments also caused step-like changes in ribosome density in individual ribosome profiles (Figure 4). Dividing the ribosome profiles for each mRNA by its corresponding RNA-Seq profiles eliminated most of these artefacts, further underlining the utility of using RNA-Seq data for Ribo-Seq analysis.

After running the change point algorithm (Muggeo and Adelfio 2011; R Development Core Team 2012), classifying the changes in density and manually inspecting the candidates for translational regulation, we found several groups of mRNAs with different types of translational regulation. Among the alternative termination targets we found four selenoprotein mRNAs (Figure 6, Table 2). Selenoproteins have an internal UGA codon that is recoded for selenocysteine insertion during translation. Previous studies have indicated that UGA recoding is not totally efficient and that selenocysteine insertion competes with premature termination (Driscoll and Copeland 2003), and therefore the presence of selenoprotein genes in this group was consistent with current understanding of selenoprotein translation. In the analysed selenoprotein ribosome profiles, ribosome density decreased at the UGA codon, but did not disappear, indicating that full size selenoproteins production and ribosome drop-off/termination occur on the same mRNA transcripts. Given the recent report that, for most selenoproteins, there is a drop in ribosome density at the UGA codon (Howard et al. 2013), we ran the algorithm specifically on all selenoproteins and found five more with the same type of change in density. *Atf4*, another discovered termination target, is a known target of uORF regulation (Vattem and Wek 2004), which was also confirmed in the original ribosome profiling study (Ingolia et al, 2011). To our knowledge, translational regulation has not yet been described for the final target identified – *Tmem55b*.

We are aware of three other studies that have used ribosome profiling to find potential TISs (Ingolia et al. 2011; Fritsch et al. 2012; Lee et al. 2012). In all studies, any site where ribosomes stalled after applying an initiation blocker were presumed to be potential TISs. However, in none of the studies was the translation associated with each of these sites systematically quantified. In our analysis we focused only on those alternative TISs in the annoatetd CDS which are associated with substantial initiation, i.e. where the ribosome density changes by at least 33 %. We found 11 genes with potential alternative initiation (Figure 7), of which five have previously been reported (Ingolia et al. 2011). A possible reason why the other six genes were missed by Ingolia et al is that harringtonine does not always halt translation at near cognate (e.g. cug, gug) TISs (Starck et al. 2008). Whilst it is possible that some of our 11 genes are false positives, it should be noted that Ingolia et al found 3,426 potential alternative TISs in the annotated CDSs. When we screened these 3,426 TISs, we discovered CPs for only 13, suggesting the vast majority of the detected TISs are associated with only limited translation.

We found a further eight mRNAs with potential ribosome drop-off, none of which were previously reported to be targets of translational regulation. The difference between these and the alternative termination mRNAs was that no stop codon was found at the positions of the change point for the latter. Analysis of the sequence surrounding the drop-off site showed enrichment for G and A nucleotides and specifically the GAA codon. A similar consensus was found for sites of ribosome pausing in the original Ribo-Seq study (Ingolia et al. 2011). It therefore seems that in at least some cases ribosome stalling can lead to ribosome drop-off in eukaryotes, as has been suggested (Buchan and Stansfield 2007).

We also found several mRNAs with high ribosome density at the 5’ (ramp), or the 3’ of the CDS (slow termination); however sequence analysis did not reveal enrichment for any sequences. Instead of resulting from translation of slow codons as it the current hypothesis, the TASEP simulations of translation have suggested that the ramp could be due to very fast initiation but this has not been confirmed by analysis of translation efficiency. Therefore, slow codons and/or mRNA tertiary structure remain the most probable causes of the ramp (Kudla et al. 2009, Tuller et al. 2010)

Although we tried to eliminate changes in density due to Ribo-Seq/RNA-Seq artefacts, a careful analysis showed that many CPs among the translation regulation candidates were a consequence of alternative splicing. Because all changes in density close to known exon/exon junctions as annotated in the UCSC Genome Browser were eliminated from the final list, the remaining CPs were strong candidates for novel isoforms. Using this approach, we found 31 genes with hitherto unknown isoforms from UCSC or Ensembl (Table S2), suggesting that our method is not only useful for analysing translational regulation, but also for uncovering novel alternative isoforms including alternative splicing. Indeed, in a very recent study, a strategy similar to ours has been developed specifically to estimate gene isoform expression with very encouraging results (Suo et al. 2013).

In conclusion, the present study demonstrates that ribosome density patterns in the CDSs are a valuable source of information in the analysis of translational regulation. The change point approach taken in the study has proven useful in detecting translational regulation in the mRNA coding region, but would easily be extendable to analysis of 5’UTR and 3’UTR regions and to detection of alternative splicing events.

## Methods

### TASEP models of protein translation

The protein translation models were developed based on the totally asymmetric simple exclusion process (TASEP) with extended particles (Lakatos and Chou 2003). In the models, an mRNA CDS with *N = 500* codons is presented by a chain of *500* sites, while a ribosome attached to the mRNA covers *L = 10* codons (Figure 1). If the first site of the mRNA is free, a new ribosome attaches at the initiation with rate *k*_*I*_. After initiation, the ribosome moves along the mRNA at a codon specific rate *k*_*Ei*_, again only if its movement is not hindered by another ribosome. When the ribosome reaches the final codon, it detaches at rate *k*_*T*_ from the mRNA together with a full-size peptide – the results of the translation process.

The dynamics of ribosome progression along the mRNA was simulated with the next reaction Gillespie algorithm (Gillespie 1977), using custom Matlab code (Supplementary File1). All simulations started with an empty mRNA, which was simulated for 1,000,000 ribosome steps. As the average half-life of a mammalian mRNA is around 9h (Schwanhausser et al. 2011) and the steady-state ribosome density is achieved in a matter of minutes, according to estimates of elongation speed of around 5aa/s (Ingolia et al. 2011), the transient lower ribosome occupation of the mRNA (first 100,000 steps) was ignored in all analyses. The ribosome profiles were determined by random sampling and averaging of ribosome position along the mRNA from step 100,000 to the end of the simulation. The protein synthesis rates were determined by dividing the total number of proteins produced by the total time that has passed from step 100,000 to the end of the simulation.

When simulating alternative initiation at a TISs other than the canonical one, the scanning model of translation initiation was assumed (Kozak 1989). Instead of the 80S ribosome coming together at the canonical TIS, the 40S skips it at rate *k*_*Alt*_ and scans downstream until a suitable TIS is found. In the simulations we assumed that the 40S scanning occurs at the same speed as elongation and that it is the same size as the full 80S ribosome, thus providing the same steric hindrance to other ribosomes. Nevertheless, since 40S ribosomes are not recorded in Ribo-Seq, we ignored them in the ribosome density calculations (Ingolia et al. 2009).

Ribosome drop-off and alternative termination were modelled as alternatives to the elongation step occurring at rate *k*_*Di*_, i.e. at any codon on the mRNA (except the final codon) the ribosome could either move to the next codon or detach from the mRNA.

### Change point analysis

In this study, we tested change point algorithms from the *changepoint* R package (http://cran.r-project.org/web/packages/changepoint/index.html) and a Bayesian change point algorithm from the *bcp* R package (http://cran.r-project.org/web/packages/bcp/index.html) (Erdman and Emerson 2008). In a preliminary run both algorithms produced very similar results, however as the *bcp* returns the estimates of both the mean and the probability of a CP for each position in a sequence (whereas *changepoint* does not), we decided to use it for the rest of the analysis. The details of the algorithm are presented in (Erdman and Emerson 2008), therefore here we only provide a broad overview and the changes we have made to the algorithm for analysis of our rather specific signals – ribosome and RNA-Seq profiles.

Ribosome and RNA-Seq profiles are represented as vectors of the number of RNA reads aligned to specific positions of the mRNA CDSs *y*_*R*_ = [*y*_*R1*_,…, *y*_*Rn*_]. The *bcp* algorithm assumes that there is an unknown partition, *ρ*, of each vector into contiguous block, such the means are equal in each block but different between neighbouring blocks. A CP is defined as the position on the mRNA *i*∊ {*1*,…, *n* - *1*} that delimits two consecutive blocks. The algorithm begins with a zero partition *ρ =* (*U*_*1*_, *U*_*2*_,…, *U*_*n*_);*U*_*i*_ = 0;*U*_*n*_ = *1* and then updates it in an MCMC scheme. In each step of the Markov chain, at each position i, a value of *U*_*i*_ is drawn from the conditional distribution of *U*_*i*_ given the data and the current partition. The transition probability, *p*, of a change at the position *i + 1,* is obtained from the following ratio presented in (Barry and Hartigan 1993):

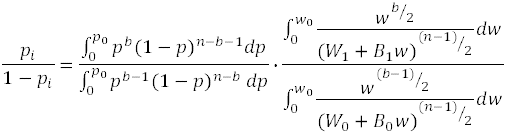

where *b* is the number of blocks, *p*_*0*_ and *w*_*0*_ are parameters used in the definition of priors that control the sensitivity of the algorithm, and *W*_*0*_,*B*_*0*_,*W*_*1*_,*B*_*1*_ are the within and between block sums of squares obtained when *U*_*i*_ = *0* and *U*_*i*_ = *1*, respectively. In this study, different values of *p*_*0*_ and *w*_*0*_ were tested, with *p*_*0*_ *=* 0.001 (the number of changes in the signal was expected to be low) and *w*_*0*_ *= 0.2* (relevant changes were expected to be of reasonable size) producing reasonable results in detecting changes in the simulated datasets (Figure S3)(Erdman and Emerson 2008). The parameters were changed to *p*_*0*_ *= 0. 1, w*_*0*_ *=* 0.02 when detecting change points after slow elongation and termination - these parameters were found to work better when the number of changes to be detected was high.

Only CPs with the following properties were included into the analysis:

- Minimum segment size of at least 10 codons – a sufficient segment size is intended to guarantee that the detected CPs are not due to random noise, but are genuine translational regulation events. In all cases, when segments were smaller, the CPs were combined into a single CP with the smallest possible increase in mean squared error between the signal and the estimated posterior means of the segments.
- The change in mean between two neighbouring segments of at least 1/3 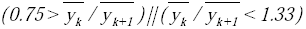. While smaller changes were detected by the algorithm, they were ignored in subsequent analysis to decrease the number of false positives.
- The confidence level of the CP had to be over 0.95.

Each detected and accepted CP was assigned an interval, which had at least 95 % probability of encompassing the true CP. The interval was defined as positions at which CL reached 2.5 % and 97.5 % of its final value, respectively.

The algorithm was first tested on 500 randomly chosen RNA-seq and ribosome profiles. The test revealed that the algorithm is sensitive to peaks in ribosome density. Because they have been studied before (Ingolia et al. 2011), we were not interested in these so called “ribosome pauses”. Therefore, we decided to scan the ribosome profiles for any peaks (3 codons or less wide) where ribosome density is at least 10 fold or greater than the mean density across the CDS and replace these peaks with the mean CDS ribosome density. We then repeated the change point analysis for the whole dataset.

### Generation of ribosome and RNA-Seq profiles for individual mRNAs

Our analysis was performed on a Ribo-Seq dataset for mouse embryonic stem cells (mESCs) (Ingolia et al. 2011) as submitted to the NCBI Gene Expression Omnibus (GEO) database (accession number GSE30839) (Barrett and Edgar 2006). From the whole set of data generated by Ingolia et al, we selected 8,933 well-expressed and translated transcripts with a CDS ribosome density > 1/nucleotide representing 4784 genes (Table S2). For each CDS, the nucleotide reads were transformed into codon reads by averaging the reads obtained from three non-overlapping consecutive nucleotides, starting from the annotated start codon and ending at the stop codon.

The average ribosome profiles and RNA-Seq profiles were generated by first normalizing all individual profiles by length. This was performed by using function *interp1* in Matlab with *‘linear’* interpolation. The profiles were then normalized against their own mean profiles values, the normalized profiles summed and the sum divided by the total number of isoforms. Finally, the ribo/RNA profiles were obtained by diving the ribosome profiles by the RNA-Seq profiles (to avoid division by zero, all zeros in RNA-Seq profiles were replaced by the minimum RNA-Seq profiles value above zero).

### Eliminating alternative splicing CPs

Based on the profile (ribosome, RNA-Seq or ribo/RNA) in which a CP was found, we categorized the CPs into seven different groups (Figure S6). Group A contains the CPs found only in ribosome profiles – they are not found in RNA-Seq profiles – which are eliminated by normalization. These are most likely to be caused by noise, by short segments of slow/fast codons or weak multiple assignments detected in ribosome profiles, but not RNA-Seq profiles (vice versa for group B). Groups D and G contain CPs found in ribosome profiles (D,G), RNA-Seq profiles (D,G) and ribo/RNA profiles (G). These are most likely due to multiple alignments/assignments. Groups C and F are most likely false positives, caused by normalization itself (C) or noise (F). We are thus left with the most likely targets for translation regulation - Group E: 1355 CPs (4 %) found in ribosome and ribo/RNA profiles, but not in RNA-Seq profiles.

This group was reduced further by eliminating all CPs that were < 3 codons away from a known exon/exon junction, and CPs that came in pairs (a rise/drop at first CP countered by a drop/rise to the same level at second CP - a pattern typical of for multiple assignments).

### Categorization of translational regulation candidates

CP properties and genomic sequence data were used to categorize CPs into groups with different translational regulation events. Increased ribosome density was taken as a sign of alternative initiation or slow termination, with decreased density indicating alternative termination, ribosome drop-off, or ramp. The position of the CP was taken into account when distinguishing between e.g. alternative initiation (more probable at 5’ ends of the CDS) and slow termination (3’ ends of the CDS). We scanned the genomic sequences surrounding the detected CPs for potential stop codons and strong initiation sequences (consensus sequence taken from Ingolia et al (Ingolia et al. 2011)) in any reading frame. We also calculated the periodicity transition score (script downloaded from http://lapti.ucc.ie/bicoding/Rscripts/PTS.R in September 2013) – a measure of how the nucleotide triplet periodicity (i.e. RNA reads being assigned either to the first, second or third nucleotide of a codon) changes in the ribosome profiles (Michel et al. 2012). If PTS > 10, we considered the possibility of a frameshift. The above data were fed into a decision tree (Figure S7) to classify the CPs automatically. Afterwards, each transcript profile was visually inspected and compared with the genomic information available in UCSC Genome Browser, especially information on all gene isoforms (Meyer et al. 2013). If the CP could be explained by a factor other than translational regulation, it was regarded as a false positive.

### Gene ontology analysis

Identified sets of genes were compared with the group of all well-expressed genes for gene ontology enrichment with the web-based tool GOrilla, using the default settings (Eden et al. 2009).

## Acknowledgements

AZ, CM, JCM, TBLK, JEH and DPS would like to thank the BBSRC for support. SNG would like to thank the Addison Wheeler Trust for support.

## References

Arava Y, Boas F, Brown P, Herschlag D. 2005. Dissecting eukaryotic translation and its control by ribosome density mapping. Nucleic Acids Res 33(8): 2421–2432.

Barrett T, Edgar R. 2006. Gene expression omnibus: Microarray data storage, submission, retrieval, and analysis. Method Enzymol 411: 352–369.

Barry D, Hartigan JA. 1993. A Bayesian-Analysis for Change Point Problems. JAm Stat Assoc 88(421): 309–319.

Bazzini A, Lee M, Giraldez A. 2012. Ribosome profiling shows that miR-430 reduces translation before causing mRNA decay in zebrafish. Science 336(6078): 233–237.

Bohnert R, Ratsch G. 2010. rQuant.web: a tool for RNA-Seq-based transcript quantitation. Nucleic Acids Res 38: W348–W351.

Brockmann R, Beyer A, Heinisch JJ, Wilhelm T. 2007. Posttranscriptional expression regulation: What determines translation rates? PLoS Comp Biol 3(3): 531–539.

Cho PF, Gamberi C, Cho-Park YA, Cho-Park IB, Lasko P, Sonenberg N. 2006. Cap-dependent translational inhibition establishes two opposing morphogen gradients in Drosophila embryos. Curr Biol 16(20): 2035–2041.

Dana A, Tuller T. 2012. Determinants of translation elongation speed and ribosomal profiling biases in mouse embryonic stem cells. PLoS Comput Biol8(11): e1002755 doi: 10.1371/journal.pcbi.1002755.

Dittmar K, Goodenbour J, Pan T. 2006. Tissue-specific differences in human transfer RNA expression. PLoS Genet 2(12): e221 doi: 10.1371/journal.pgen.0020221.

Driscoll D, Copeland P. 2003. Mechanism and regulation of selenoprotein synthesis. Annu Rev Nutr 23: 17–40.

Eden E, Navon R, Steinfeld I, Lipson D, Yakhini Z. 2009. GOrilla: a tool for discovery and visualization of enriched GO terms in ranked gene lists. BMC Bioinformatics 10 doi: 10.1186/1471-2105-10-48.

Erdman C, Emerson JW. 2008. A fast Bayesian change point analysis for the segmentation of microarray data. Bioinformatics 24(19): 2143–2148.

Fritsch C, Herrmann A, Nothnagel M, Szafranski K, Huse K, Schumann F, Schreiber S, Platzer M, Krawczak M, Hampe J et al. 2012. Genome-wide search for novel human uORFs and N-terminal protein extensions using ribosomal footprinting. Genome Res 22(11):2208–2218.

Gillespie DT. 1977. Exact Stochastic Simulation of Coupled Chemical-Reactions. J Phys Chem-Us 81(25): 2340–2361.

Guo H, Ingolia NT, Weissman JS, Bartel DP. 2010. Mammalian microRNAs predominantly act to decrease target mRNA levels. Nature 466(7308): 835–840.

Hansen KD, Brenner SE, Dudoit S. 2010. Biases in Illumina transcriptome sequencing caused by random hexamer priming. Nucleic Acids Res 38(12): e131 doi: 10.1093/nar/gkq224.

Hesketh J. 2008. Nutrigenomics and selenium: Gene expression patterns, physiological. targets, and genetics. Annu Rev Nutr 28: 157–177.

Howard MT, Carlson BA, Anderson CB, Hatfield DL. 2013. Translational Redefinition of UGA Codons Is Regulated by Selenium Availability. J BiolChem 288(27): 19401–19413.

Ingolia N, Lareau L, Weissman J. 2011. Ribosome profiling of mouse embryonic stem cells reveals the complexity and dynamics of mammalian proteomes. Cell 147(4): 789–802.

Ingolia NT, Brar GA, Rouskin S, McGeachy AM, Weissman JS. 2012. The ribosome profiling strategy for monitoring translation in vivo by deep sequencing of ribosome-protected mRNA fragments. Nat Protoc 7(8): 1534–1550.

Ingolia NT, Ghaemmaghami S, Newman JR, Weissman JS. 2009. Genome-wide analysis in vivo of translation with nucleotide resolution using ribosome profiling. Science 324(5924): 218–223.

Ingolia NT. 2014. Ribosome profiling: new views of translation, from single codons to genome scale. Nat Rev Genet: published online 28 January doi: 10.1038/nrg3645.

Karolchik D, Hinrichs AS, Furey TS, Roskin KM, Sugnet CW, Haussler D, Kent WJ. 2004. The UCSC Table Browser data retrieval tool. Nucleic Acids Res 32: D493–D496.

Kozak M. 1989. The Scanning Model for Translation – an Update. J Cell Biol 108(2): 229–241.

Kudla G, Murray A, Tollervey D, Plotkin J. 2009. Coding-sequence determinants of gene expression in Escherichia coli. Science 324(5924): 255–258.

Kurland CG. 1992. Translational Accuracy and the Fitness of Bacteria. Annu Rev Genet 26: 29–50.

Lakatos G, Chou T. 2003. Totally asymmetric exclusion processes with particles of arbitrary size. J Phys A Math Gen 36: 2027–2041.

Lee S, Liu B, Lee S, Huang S-X, Shen B, Qian S-B. 2012. Global mapping of translation initiation sites in mammalian cells at single-nucleotide resolution. PNAS 109(37): E2424–E2432.

Leprivier G, Remke M, Rotblat B, Dubuc A, Mateo AR, Kool M, Agnihotri S, El-Naggar A, Yu B, Somasekharan SP et al. 2013. The eEF2 kinase confers resistance to nutrient deprivation by blocking translation elongation. Cell 153(5): 1064–1079.

Li GW, Oh E, Weissman JS. 2012. The anti-Shine-Dalgarno sequence drives translational pausing and codon choice in bacteria. Nature 484(7395): 538–541.

Liu B, Han Y, Qian SB. 2013. Cotranslational response to proteotoxic stress by elongation pausing of ribosomes. Mol Cell 49(3): 453–463.

Meyer LR, Zweig AS, Hinrichs AS, Karolchik D, Kuhn RM, Wong M, Sloan CA, Rosenbloom KR, Roe G, Rhead B et al. 2013. The UCSC Genome Browser database: extensions and updates 2013. Nucleic Acids Res 41(D1): D64–D69.

Michel A, Choudhury K, Firth A, Ingolia N, Atkins J, Baranov P. 2012. Observation of dually decoded regions of the human genome using ribosome profiling data. Genome Res 22(11): 2219–2229.

Muggeo VMR, Adelfio G. 2011. Efficient change point detection for genomic sequences of continuous measurements. Bioinformatics 27(2): 161–166.

R Development Core Team (2008). R: A language and environment for statistical computing. R Foundation for Statistical Computing, Vienna, Austria. ISBN 3-900051-07-0, URL http://www.R-project.org.

Richter JD, Sonenberg N. 2005. Regulation of cap-dependent translation by eIF4E inhibitory proteins. Nature 433(7025): 477–480.

Roberts A, Trapnell C, Donaghey J, Rinn JL, Pachter L. 2011. Improving RNA-Seq expression estimates by correcting for fragment bias. Genome Biol 12(3): R22 doi: 10.1186/gb-2011-12-3-r22.

Rogers AN, Chen D, McColl G, Czerwieniec G, Felkey K, Gibson BW, Hubbard A, Melov S, Lithgow GJ, Kapahi P. 2011. Life span extension via eIF4G inhibition is mediated by posttranscriptional remodeling of stress response gene expression in C. elegans. Cell Metab 14(1): 55–66.

Schwanhausser B, Busse D, Li N, Dittmar G, Schuchhardt J, Wolf J, Chen W, Selbach M. 2011. Global quantification of mammalian gene expression control. Nature 473(7347): 337–342.

Shaw L, Zia R, Lee K. 2003. Totally asymmetric exclusion process with extended objects: a model for protein synthesis. Phys Rev E 68(2): 21910 doi: 10.1103/PhysRevE.68.021910

Sonenberg N, Hinnebusch A. 2009. Regulation of translation initiation in eukaryotes: mechanisms and biological targets. Cell 136(4): 731–745.

Stadler M, Fire A. 2011. Wobble base-pairing slows in vivo translation elongation in metazoans. RNA 17(12): 2063–2073.

Starck SR, Ow YK, Jiang V, Tokuyama M, Rivera M, Qi X, Roberts RW, Shastri N. 2008. A Distinct Translation Initiation Mechanism Generates Cryptic Peptides for Immune Surveillance. PloS ONE 3(10): e3460 doi: 10.1371/journal.pone.0003460.

Suo C, Calza S, Salim A, Pawitan Y. 2013. Joint estimation of isoform expression and isoform-specific read distribution using multisample RNA-Seq data. *Bioinformatics* ahead of print doi: 10.1093/bioinformatics/btt704.

Sutton MA, Taylor AM, Ito HT, Pham A, Schuman EM. 2007. Postsynaptic decoding of neural activity: eEF2 as a biochemical sensor coupling miniature synaptic transmission to local protein synthesis. Neuron 55(4): 648–661.

Tian Q, Stepaniants S, Mao M, Weng L, Feetham M, Doyle M, Yi E, Dai H, Thorsson V, Eng J et al. 2004. Integrated genomic and proteomic analyses of gene expression in Mammalian cells. Mol Cell Proteomics 3(10): 960–969.

Tuller T, Carmi A, Vestsigian K, Navon S, Dorfan Y, Zaborske J, Pan T, Dahan O, Furman I, Pilpel Y. 2010. An evolutionarily conserved mechanism for controlling the efficiency of protein translation. Cell 141(2): 344–354.

Vattem KM, Wek RC. Reinitiation involving upstream ORFs regulates ATF4 mRNA translation in mammalian cells. PNAS 101(31): 11269–11274.

Vogel C, Abreu R, Ko D, Le S-Y, Shapiro B, Burns S, Sandhu D, Boutz D, Marcotte E, Penalva L. 2010. Sequence signatures and mRNA concentration can explain two-thirds of protein abundance variation in a human cell line. Mol Syst Biol 6: 400 doi: 10.1038/msb.2010.59.

